# Prediction of antibiotic resistant strains of bacteria from their beta-lactamases protein

**DOI:** 10.1101/2021.06.26.450028

**Authors:** Lubna Maryam, Anjali Dhall, Sumeet Patiyal, Salman Sadullah Usmani, Neelam Sharma, Gajendra Pal Singh Raghava

## Abstract

Number of beta-lactamase variants have ability to deactivate ceftazidime antibiotic, which is the most commonly used antibiotic for treating infection by Gram-negative bacteria. In this study an attempt has been made to develop a method that can predict ceftazidime resistant strains of bacteria from amino acid sequence of beta-lactamases. We obtained beta-lactamases proteins from the β-lactamase database, corresponding to 87 ceftazidime-sensitive and 112 ceftazidime-resistant bacterial strains. All models developed in this study were trained, tested, and evaluated on a dataset of 199 beta-lactamases proteins. We generate 9149 features for beta-lactamases using Pfeature and select relevant features using different algorithms in scikit-learn package. A wide range of machine learning techniques (like KNN, DT, RF, GNB, LR, SVC, XGB) has been used to develop prediction models. Our random forest-based model achieved maximum performance with AUROC of 0.80 on training dataset and 0.79 on the validation dataset. The study also revealed that ceftazidime-resistant beta-lactamases have amino acids with non-polar side chains in abundance. In contrast, ceftazidime-sensitive beta-lactamases have amino acids with polar side chains and charged entities in abundance. Finally, we developed a webserver “ABCRpred”, for the scientific community working in the era of antibiotic resistance to predict the antibiotic resistance/susceptibility of beta-lactamase protein sequences. The server is freely available at (http://webs.iiitd.edu.in/raghava/abcrpred/).

**Key Points:**

- Ceftazidime is commonly used to treat infection caused by Gram-negative bacteria.
- Beta-lactamase is responsible for lysing ceftazidime, make it resistant to bacteria.
- Comparison of resistant and sensitive variants of beta-lactamase.
- Classification of sensitive and resistant strain of bacteria based on beta-lactamase.
- Prediction models have been developed using different machine learning techniques.

## Introduction

Antimicrobial resistance (AMR) is the ability of bacteria to resist the effect of antibiotics that are administered during infection (Figure 1). In 2020, WHO declared AMR as one of the world’s top 10 public health threats. There is a number of reasons for drug resistance that include overuse of antibiotics and the emerging strain of bacteria. Moreover, the alarming spread of multi-drug resistant bacteria (MDR) continues to lurk our capability to treat common infections (e.g., sepsis, sexually transmitted infections, urinary tract infections, diarrhea). There are a number of mechanisms adopted by the bacteria to evade killing by antimicrobial molecules that include, production of antibiotic lysing enzymes (e.g., Beta-lactamases), lowering the permeability of cell membrane, and modification of the antibiotics binding site [1]. Beta-lactam antibiotics are the most prescribed antibiotics to fight broad spectrum infections, i.e., 65% of the total antibiotics in the market [2]. These antibiotics have four membered beta-lactam rings in their molecular structure, which are destroyed by beta-lactamases [3]. Thus, beta-lactamases are responsible for multi-drug resistance against beta-lactam antibiotics [4]. The number of beta-lactamases is continuously growing, around 7166 beta-lactamases have been already identified [5]. There are only a few variants of beta-lactamases on which beta-lactam antibiotics are working (sensitive). Resistant beta-lactamase genes that are spread via diffusion of mobile genetic elements, spread of epidemic plasmids, dispersion of specific clones and horizontal gene transfer [6]. Therefore, there is an urgent need to develop prediction models that can discriminate antibiotics sensitive and resistant variants of beta-lactamases.

**Figure 1:**
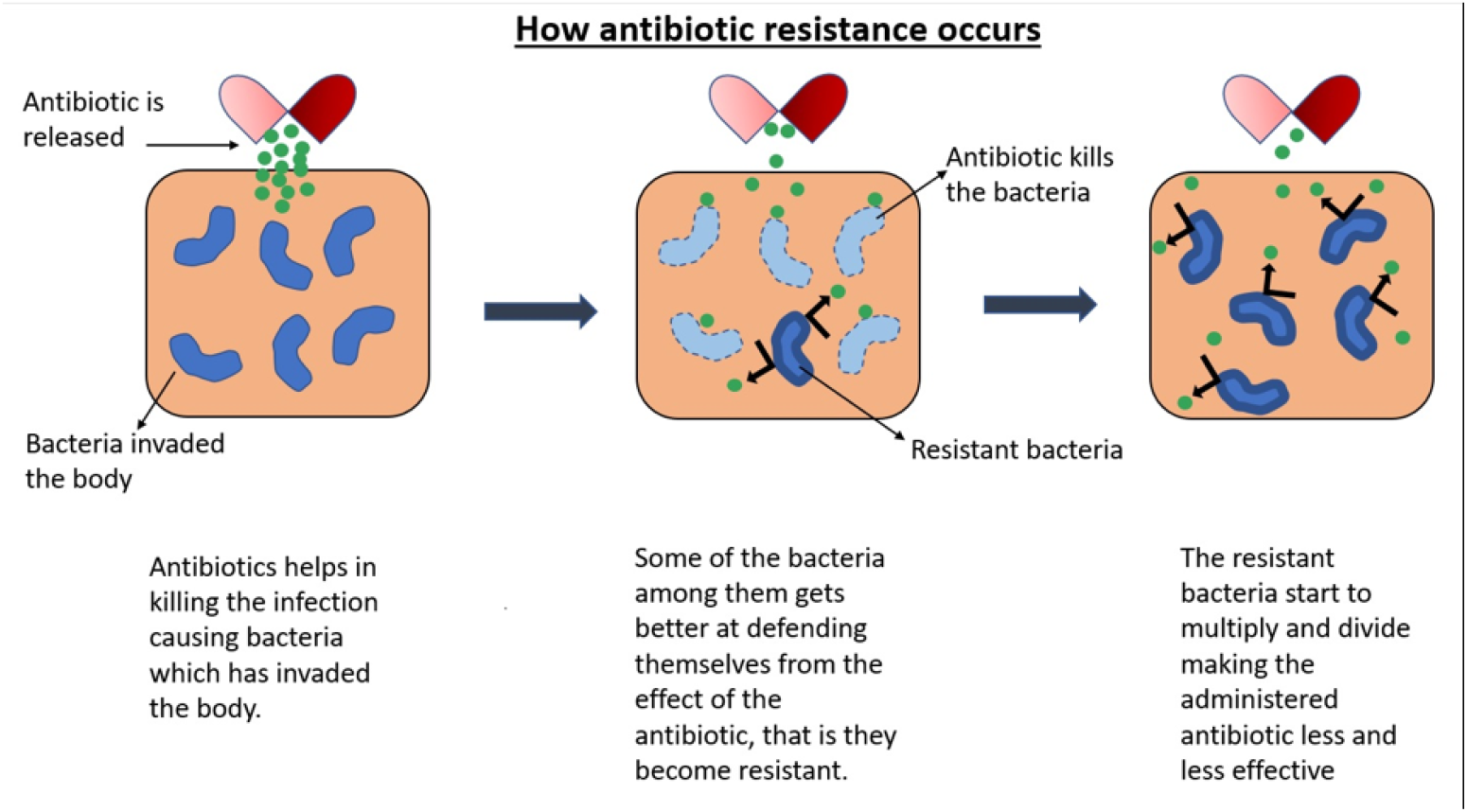
Pictorial representation of how antibiotic resistance occurs.

The standard technique to test the resistance and susceptibility of a strain to a particular antibiotic is the disc diffusion test. This method is reliable and reproducible, but it is time consuming and labour-intensive. Thus, there is a need to develop computational methods that can predict antibiotic resistant strains of bacteria. Due to advancements in the next-generation sequencing (NGS) technologies, it is in routine to sequence a gene or whole genome of bacteria or metagenome. Several repositories have been already developed to maintain the information regarding the genes, mutations, genomes and metagenomes of bacterial strains [7]. This information has been used to develop methods for predicting drug resistant strains of bacteria. These methods are mainly based on identification of antibiotic resistant gene, mutation, whole genome and metagenome [8–11]. Almost all the existing tools are generic in nature, where these methods predict whether a bacterial strain is resistant to all antibiotics. In other words, these methods predict multi-antibiotics resistant bacteria. In the past, a large number of antibiotics have been discovered to kill bacteria by a different mechanism. It is possible that bacterial strain is only resistant to a particular antibiotic or class of antibiotics but sensitive to other class of antibiotics. Thus, it is important to develop a method that can predict antibiotic-specific sensitive or resistant bacterial strains; similar to personalized medicine [12–14]. This is very important to manage treatment of bacterial infection using a particular antibiotic which is sensitive to bacteria responsible for a given infection. In simple worlds there is need to treat a bacterial infection using strain-specific antibiotics which is similar to personalize drugs. Best of our knowledge, there is no computational tool that can predict whether a bacterial strain is sensitive or resistant to a antibiotics. In this study, we first time made an attempt to develop method for antibiotic ceftazidime that belongs to beta-lactam group. We selected ceftazidime because it is routinely used for treatment of wide range of bacterial infections like meningitis, sepsis, joint infection, urinary tract infection. In addition, ceftazidime has been tested clinically on a number of bacterial strains where MIC have been determined. In order to identify sensitive and resistant strain from MIC values, European Committee on Antimicrobial Susceptibility Testing (2020) proposed that the Enterobacterales are susceptible to ceftazidime when its concentration is less than or equal to 1 mg/ml and resistant when concentration is greater than 4 mg/ml. It is well known fact that beta-lactamases are responsible for lysing ceftazidime or causing resistance. Thus, we have designed a model for predicting beta-lactamase variants that make ceftazidime sensitive or resistant to a bacterial strain. We used state of the arts techniques mainly based on machine learning techniques to develop prediction models [15]. This will help in predicting ceftazidime resistance/susceptibility towards beta-lactamase carrying bacterial species that could emerge in the near future. The platform will provide vista to find out the beta-lactamases strains which are sensitive to ceftazidime antibiotic.

## Methods and Material

### Dataset Collection

The main dataset was collected from the β-lactamases database [16]. It incorporates 2383 Minimum Inhibitory Concentration (MIC) values (in the presence and absence of beta-lactamase genes) of 980 beta-lactamases (with their protein sequences) with different antibiotics and the fold change in MIC values [16].The database comprises experimentally validated 21 different types of antibiotics corresponding to class-A, B, C, D beta-lactamase proteins. In this study, we have considered the ceftazidime antibiotic dataset with different β-lactamase protein sequences. Our final dataset included 199 β-lactamase protein sequences. Further, we set a cutoff on MIC values of ceftazidime with β-lactamase proteins. The proteins having (MIC value <=4) were considered as antibiotic susceptible/sensitive proteins, and proteins having (MIC value >4) were taken as antibiotic resistant ones [17,18]. Finally, we got 87 antibiotic-sensitive and 112 antibiotic-resistant unique proteins, referred to as positive and negative dataset, respectively. Moreover, we have also collected 22 ceftazidime resistance beta-lactamase protein sequences from Resistance Gene Identifier (RGI) database for external validation [19].

### Generation of Features

To generate a wide range of features from protein sequences, we have used Pfeature [20]. In this study, we have used the standalone package of the Pfeature tool to compute thousands of protein/peptide features. This tool also calculates the structural and functional properties of protein sequences. We have generated 9149 composition-based features/descriptors using the composition-based feature module of the Pfeature package. It incorporates 15 different type of descriptors such as Amino acid composition (AAC), Dipeptide composition (DPC), Tripeptide composition (TPC), Atomic and bond composition (ABC), Residue repeat Information (RRI), Distance distribution of residue (DDOR), Shannon-entropy of protein (SE), Shannon entropy of all amino acids (SER), Shannon entropy of physicochemical property (SEP), Conjoint triad calculation of the descriptors (CTD), Composition-enhanced transition distribution (CeTD), Pseudo amino acid composition (PAAC), Amphiphilic pseudo amino acid composition (APAAC), Quasi-sequence order (QSO) and Sequence order coupling number (SOCN).

### Pre-processing and Feature selection

The biggest challenge is to find out the most important features/descriptors which can classify the two classes more accurately. The standardization or scaling of the dataset is the most common requirement for the machine learning techniques. In the current study, to standardize the dataset, we used MinMaxScaler using the sklearn pre-processing package. This scaling function converts the given values into a minimum and maximum range. After the pre-processing step, we identified the best set of features from a huge dimension vector. For determining the best features several dimension reduction methods are currently available. We have used standard feature selection methods in which firstly we removed all low variance features using the variance threshold method of the scikit-learn package [21]. It removes all zero-variance features, so we were left with 275 features. Then, we applied the SVC-L1 feature selection method for the selection of important set of features [21]. This method is based on the support vector classifier (SVC) with linear kernel, penalized with L1 regularization. SVC-L1 method was performed on earlier deduced 275 features which provided 33 features. Further we ranked the features based on their performance, using feature selector tool. We developed our final machine learning models on selected 10, 20, and 33 features.

### Machine Learning

In the present study, we have implemented several machine learning techniques to classify ceftazidime antibiotic-resistant and sensitive/non-resistant proteins. We incorporated K-nearest neighbors (KNNs), Decision tree (DT), Random Forest (RF), Gaussian Naive Bayes (GNB), Logistic Regression (LR) and Support Vector Classifier (SVC) and XGBoost (XGB) classification methods in the study (ref). These techniques are based on different algorithm such as, KNN is a simple and supervised machine learning algorithm. It assumes the similarity between the new data and the available data and put the new data into the category that is most similar to the available categories [22]. DT is a tree-structured classifier based on non-parametric machine learning models, which uses a decision tree as a model to go from observations about a data to conclusions about new data. RF classification method uses ensemble-based techniques which uses several decision trees for the training and prediction of the outcome [23], GNB (Gaussian Naïve Bayes) are a group of supervised classification algorithms based on Bayes theorem which uses probabilistic approach for the classification. LR is a statistical model that measures the relationship between the categorial dependent variable and one or more independent variable by guessing the likelihoods using a logistic function [24]. SVC get the best fit of the data provided, the features can then be fed to see what the predicted class is. XGB uses an iterative approach for the classification. It is a decision tree-based ensemble machine learning technique that uses an approach where new models are created that predict the errors of prior models and then added together to make the final prediction. All these techniques were executed using python-library scikit-learn [21].

### Evaluation Techniques

In order to evaluate the classification models, we have used five-fold cross-validation (CV) and external validation method. For the training, testing, and evaluation, the dataset was divided into 80:20 ratio. We have used the standard criterion for the evaluation, in which 80% of the data was used for training and 20% was used for external validation [25]. In 5-fold CV, 80% of the data was divided into five equal portions/folds, one-fold was used for testing, and four folds was used for the training purpose. A similar process was repeated five times, in which each portion/fold was utilized for internal training and testing. Further, we checked the performance of machine learning models on external dataset. In this study we have used well established evaluation parameters [26]. It incorporates threshold-dependent and independent parameters. We measured threshold-dependent parameters like sensitivity (Sens), Specificity (Spec), Accuracy (Acc) and Matthews correlation coefficient (MCC) with the help of following equations. The standard threshold-independent parameter is Area Under the Receiver Operating Characteristic (AUROC) curve [27–29] which was computed to estimate the performance of different modes.

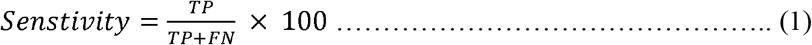

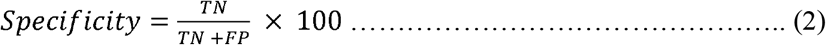

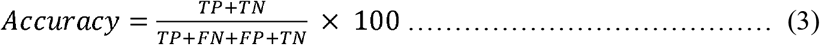

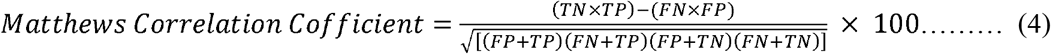

The measurements obtained from the above parameters are expressed in terms of TP=True Positive, FP=False Positive, TN=True Negative, FN=False Negative.

## Results

We have used 87 beta-lactamases proteins that are ceftazidime sensitive, having MIC values less than or equal to 4 and 112 ceftazidime resistant beta-lactamase protein sequences with MIC values greater than 4. All analysis and model development have been done on the above dataset.

### Analysis based on the amino acid composition

We have analyzed the average amino acid composition for each residue for ceftazidime resistant and sensitive beta-lactamase sequences and found out that residues such as A, G, L, P, and R, is higher in ceftazidime resistant beta-lactamase sequences as compared to sensitive sequences. Whereas, in the case of ceftazidime sensitive sequences, D, I, K, N, T, and Y residues are higher, as shown in Figure 2.

**Figure 2:**
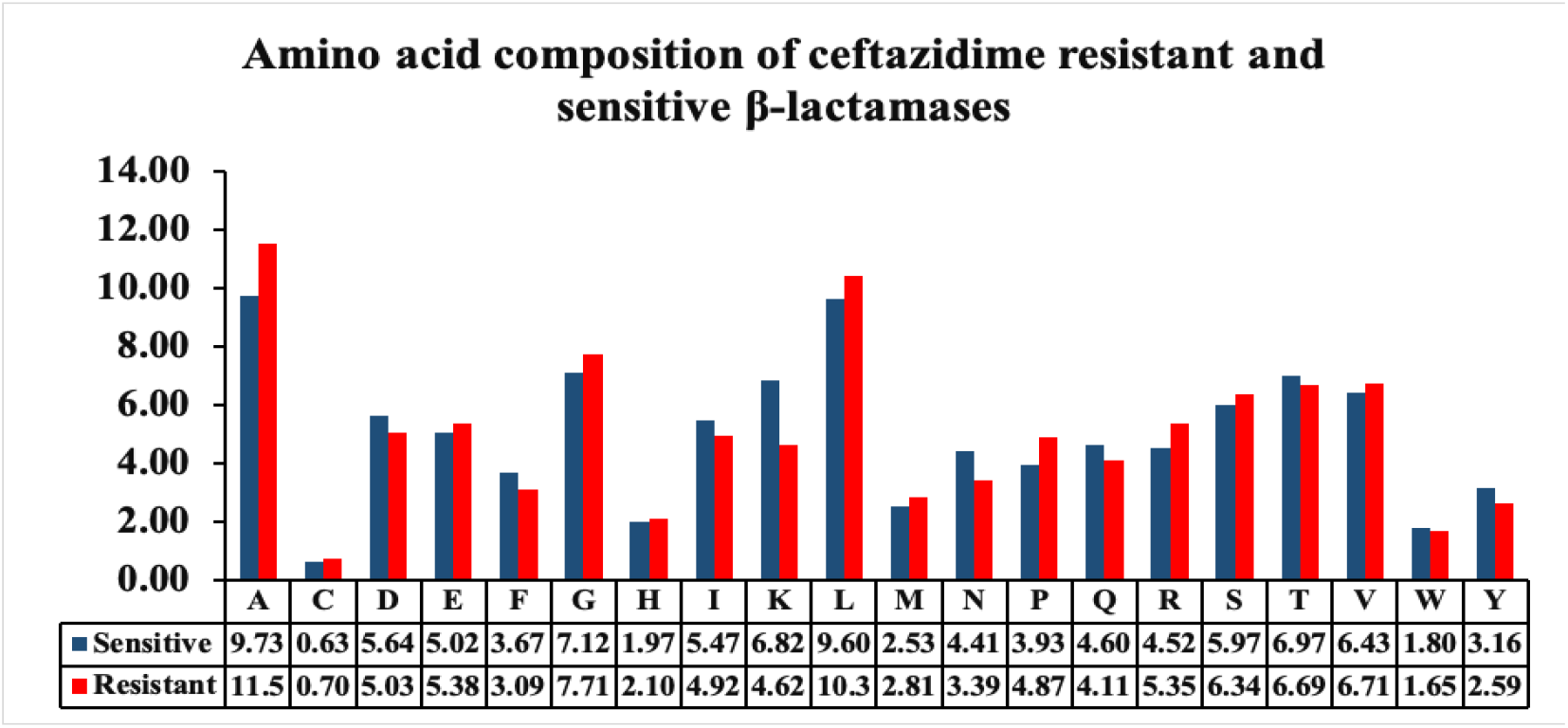
Average amino acid composition of each amino acid residues for ceftazidime-resistant and ceftazidime-sensitive beta-lactamases.

### Predictions based on machine-learning models

We have implemented various machine learning classifiers such as KNN, DT, RF, GNB, LR, SVC and XGB to develop the prediction model to classify the sequences of ceftazidime resistant and sensitive beta-lactamases. We have calculated each protein sequence’s features using the composition-based module of Pfeature, which resulted in 9149 features. On applying the feature selection method using the support vector classifier with L1 regularization, we were left with 33 most relevant features. We have ranked these 33 features using feature selector python package and generated prediction models for the top 10, 20, and 33 features. For top 10 features, RF has obtained balanced results with AUROC of 0.78 with MCC of 0.44 for training dataset, whereas AUROC and MCC for the validation dataset are 0.76 and 0.49, respectively. Performance for all the implemented classifier is exhibited in Table 2. To understand the difference between the positive and negative datasets, we calculated the average values of the top-10 features of ceftazidime-sensitive and ceftazidime-resistant beta-lactamases as represented in Table 1.

**Table 1:** Brief description of top 10 features and their average values in ceftazidime-sensitive and ceftazidime-resistant beta-lactamases.

| Name of features | Description of features | #Average Value-1 | #Average Value-2 |
| --- | --- | --- | --- |
| total_bonds | Bond composition of peptide | 4980.345 | 5353.161 |
| hydrogen_bonds | Bond composition of peptide | 2747.425 | 2958.643 |
| single_bond | Bond composition of peptide | 4544.747 | 4890.884 |
| R_ddor | Distance distribution of Arginine | 47.46805 | 36.44679 |
| Y_ddor | Distance distribution of Tyrosine | 55.61483 | 69.62223 |
| Grantham_gap1 | Quasi sequence order of peptide | 9422.904 | 9228.332 |
| Grantham_gap3 | Quasi sequence order of peptide | 9815.743 | 9486.559 |
| CeTD_33_SA | Number of transitions taking place from group 1 residues to group 2 residues for solvent accessibility attribute | 65.29885 | 53.53571 |
| CeTD_22_PC | Number of transitions taking place from group 2 residues to group 2 residues for polarizability attribute | 66.2069 | 55.08036 |
| CeTD_33_PO | Number of transitions taking place from group 3 residues to group 3 residues for polarity attribute | 67.4023 | 53.71429 |

**#Average Value-1:** average values of ceftazidime-resistance beta-lactamases; **# Average Value-2:** average values of ceftazidime-sensitive beta-lactamases.

**Table 2:** Performance of various classifiers using top 10 features.

| Classifier | Training Dataset | Validation dataset |
| --- | --- | --- |

|  | **Sens** | **Spec** | **Acc** | **AUROC** | **MCC** | **Sens** | **Spec** | **Acc** | **AUROC** | **MCC** |
| --- | --- | --- | --- | --- | --- | --- | --- | --- | --- | --- |
| KNN | 83.33 | 67.21 | 76.26 | 0.83 | 0.51 | 76.47 | 57.69 | 68.33 | 0.72 | 0.35 |
| DT | 60.26 | 72.13 | 65.47 | 0.72 | 0.32 | 76.47 | 65.38 | 71.67 | 0.72 | 0.42 |
| RF | 76.92 | 67.21 | 72.66 | 0.78 | 0.44 | 76.47 | 73.08 | 75.00 | 0.76 | 0.49 |
| GNB | 55.13 | 77.05 | 64.75 | 0.71 | 0.32 | 23.53 | 80.77 | 48.33 | 0.64 | 0.05 |
| LR | 69.23 | 68.85 | 69.06 | 0.73 | 0.38 | 61.76 | 65.38 | 63.33 | 0.67 | 0.27 |
| SVC | 76.92 | 78.69 | 77.70 | 0.78 | 0.55 | 70.59 | 73.08 | 71.67 | 0.74 | 0.43 |
| XGB | 78.21 | 65.57 | 72.66 | 0.75 | 0.44 | 76.47 | 65.38 | 71.67 | 0.78 | 0.42 |

Similarly, RF model developed using top 20 features performed best among all the classifiers, with AUROC of 0.79 and MCC of 0.48 on training dataset, and AUROC 0.76 and MCC 0.4 on validation dataset. Performance using other classifiers is given in Table 3.

**Table 3:** Performance of various classifiers using top 20 features.

| **Classifier** | **Training Dataset** |  |  |  |  | **Validation dataset** |  |  |  |  |
| --- | --- | --- | --- | --- | --- | --- | --- | --- | --- | --- |
|  | **Sens** | **Spec** | **Acc** | **AUROC** | **MCC** | **Sens** | **Spec** | **Acc** | **AUROC** | **MCC** |
| KNN | 78.21 | 68.85 | 74.10 | 0.82 | 0.47 | 73.53 | 57.69 | 66.67 | 0.71 | 0.32 |
| DT | 61.54 | 62.30 | 61.87 | 0.67 | 0.24 | 70.59 | 61.54 | 66.67 | 0.68 | 0.32 |
| RF | 74.36 | 73.77 | 74.10 | 0.79 | 0.48 | 67.65 | 73.08 | 70.00 | 0.76 | 0.40 |
| GNB | 53.85 | 77.05 | 64.03 | 0.70 | 0.31 | 29.41 | 73.08 | 48.33 | 0.63 | 0.03 |
| LR | 70.51 | 70.49 | 70.50 | 0.77 | 0.41 | 76.47 | 65.38 | 71.67 | 0.69 | 0.42 |
| SVC | 56.41 | 85.25 | 69.06 | 0.75 | 0.43 | 64.71 | 84.62 | 73.33 | 0.74 | 0.49 |
| XGB | 75.64 | 63.93 | 70.50 | 0.72 | 0.40 | 70.59 | 65.38 | 68.33 | 0.71 | 0.36 |

For all 33 features, RF obtained the maximum AUROC of 0.80 with 0.48 MCC on the training dataset, and AUROC of 0.79 with MCC of 0.46 on the validation dataset. We have reported performance for all classifiers using 33 features in the Table 4.

**Table 4:** Performance of various classifiers using 33 selected features on training and validation datasets.

| **Classifier** | **Training Dataset** |  |  |  |  | **Validation dataset** |  |  |  |  |
| --- | --- | --- | --- | --- | --- | --- | --- | --- | --- | --- |
|  | **Sens** | **Spec** | **Acc** | **AUROC** | **MCC** | **Sens** | **Spec** | **Acc** | **AUROC** | **MCC** |
| KNN | 74.36 | 77.05 | 75.54 | 0.81 | 0.51 | 73.53 | 57.69 | 66.67 | 0.74 | 0.32 |
| DT | 65.38 | 70.49 | 67.63 | 0.72 | 0.36 | 79.41 | 69.23 | 75.00 | 0.73 | 0.49 |
| RF | 74.35 | 73.77 | 74.10 | 0.80 | 0.48 | 73.53 | 73.08 | 73.33 | 0.79 | 0.46 |
| GNB | 56.41 | 72.13 | 63.31 | 0.67 | 0.29 | 29.41 | 73.08 | 48.33 | 0.64 | 0.03 |
| LR | 74.36 | 65.57 | 70.50 | 0.77 | 0.40 | 76.47 | 69.23 | 73.33 | 0.78 | 0.46 |
| SVC | 57.69 | 88.52 | 71.22 | 0.74 | 0.47 | 50.00 | 88.46 | 66.67 | 0.74 | 0.40 |
| XGB | 74.36 | 73.77 | 74.10 | 0.77 | 0.48 | 76.47 | 76.92 | 76.67 | 0.79 | 0.53 |

In order to check the robustness of our final model, we have downloaded 22 ceftazidime resistant protein sequences from Resistance Gene Identifier (RGI) database and checked the performance by implementing random forest based model developed on top 33 features. 19 out of 22 sequences were giving the correct result, with AUROC of 0.81 and MCC of 0.50 on training dataset, and AUROC 0.79 and MCC 0.71 on validation dataset.

### Webserver implementation

We have developed a webserver named ABCRpred (https://webs.iiitd.edu.in/raghava/abcrpred/) using Random Forest based machine learning approach to serve the scientific world. Since, we wanted to identify sensitive strains of beta-lactamases therefore we developed this method to discriminate between antibiotic resistant and sensitive variants of beta-lactamase strains. 87 antibiotic-sensitive and 112 antibiotic-resistant beta-lactamases protein sequences data were used for training and testing, while building the webserver. The complete architecture of ABCRpred is shown in figure 3.

**Figure 3:**
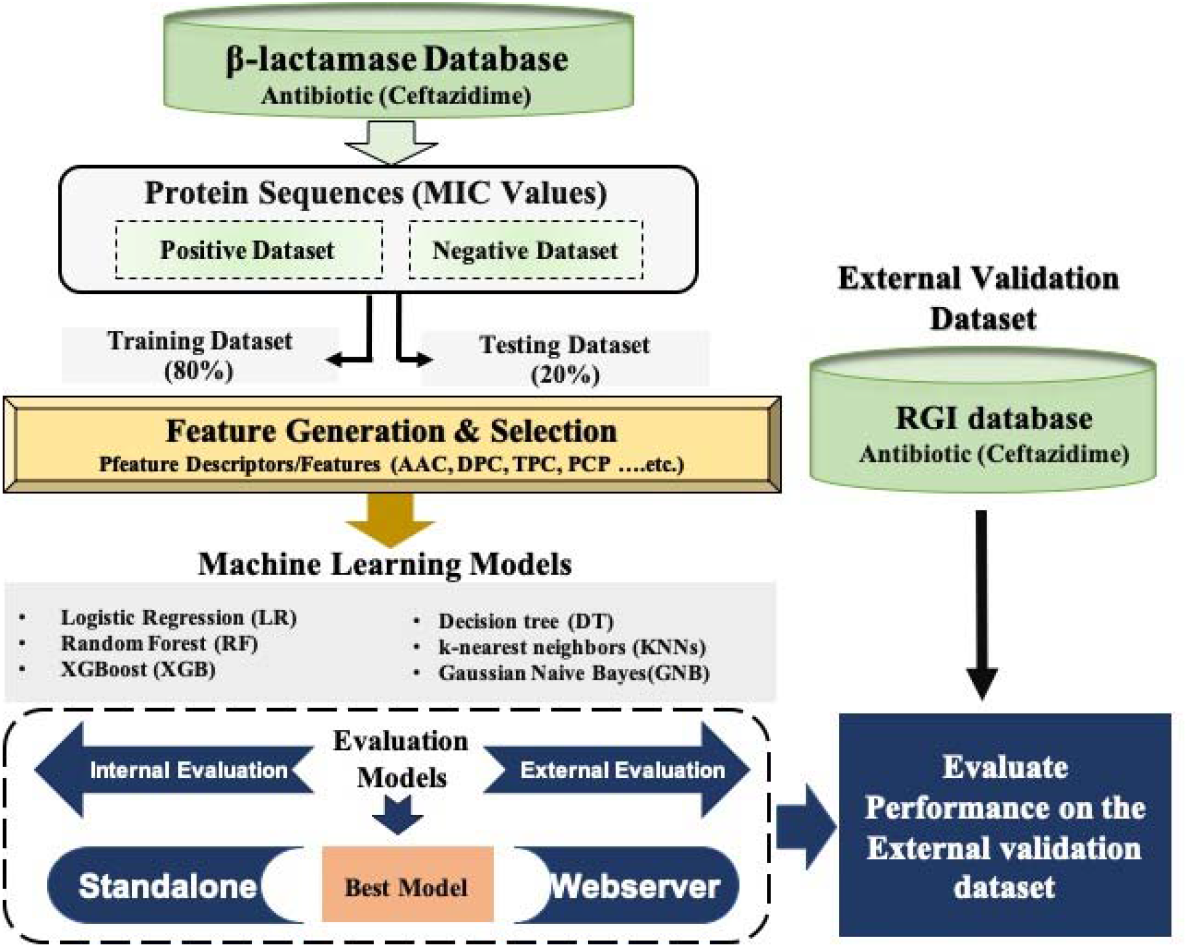
Overall ABCRpred architecture that shows process of creating datasets, features selection, model development and process of model evaluation.

The ‘Predict’ page on the webserver has been developed to predict resistance/susceptibility of any new beta-lactamase protein sequence towards ceftazidime antibiotic. The page enables the users to enter the sequence in FASTA format or upload the file with multiple peptide sequences. User is required to set a random forest threshold and select physicochemical properties as per their requirement. Prediction of each sequence will be carried out according to the selected model. After submitting the input, the output file contains various columns of sequence ID, random forest score, prediction outcome whether the input sequence is resistant or susceptible and the result of selected physicochemical properties. The standalone package (https://webs.iiitd.edu.in/raghava/abcrpred/stand.php) has also been incorporated in the webserver to let the users predict the resistance/susceptibility profile of protein sequences even in the absence of the internet. The standalone version incorporated our best models and can work on Linux or Unix operating systems.

## Discussion

The beta-lactam antibiotics are regarded as the drug of choice for the treatment of severe infections caused by *Enterobacteriaceae*. Most of the beta-lactam antibiotics face resistance against beta-lactamases carrying bacteria. Moreover, exposure of beta-lactamase carrying bacterial strains to multitude of beta-lactams has induced active continuous production and mutation of beta-lactamases expanding their activity even against the newly developed beta-lactam antibiotics. In this study we used MIC data of ceftazidime (a beta-lactam antibiotic) against beta-lactamase carrying bacteria for building a prediction model to predict resistance and susceptibility of any newly emerged variant of beta-lactamase carrying bacterial strain. A total of 199 experimental MIC data was collected from a comprehensive database of beta-lactamase enzymes called as β-lactamase Database [16]. Our data of ceftazidime MIC against various beta-lactamase carrying bacterial strains was divided into two sets. One set have MIC values greater than 4 referred to as ceftazidime-resistant strain; other set have MIC values less than or equal to 4 called as ceftazidime-sensitive strains. We obtained beta-lactamase corresponding to each strain; a beta-lactamase corresponding to resistant strain is called resistant beta-lactamase and a beta-lactamase corresponding to sensitive strain is called sensitive beta-lactamase. Our final dataset have sensitive and resistant variants of beta-lactamase.

Amino acid composition analysis revealed that certain residues like Alanine, Glycine, Leucine, Proline and Arginine are more frequent in ceftazidime resistant beta-lactamases as compared to ceftazidime sensitive ones. Similarly, in case of ceftazidime sensitive beta-lactamases the residues like Aspartic acid, Isoleucine, Lysine, Asparagine, Threonine and Tyrosine are more in abundance in comparison to resistant beta-lactamases. From these findings it can be inferred that in ceftazidime-resistant beta-lactamases, amino acids with non-polar side chains predominates. No wonder this gives these resistant beta-lactamases extra stability making it hard for ceftazidime to inhibit their activity. In case of ceftazidime-sensitive beta-lactamases, amino acids with polar side chains predominates. Moreover, amino acid with charged entities is more in number in this case. This makes these proteins quite unstable and prone to attack by the antibiotic.

In this study, Pfeature software has been used to compute different types of descriptors that includes amino acid composition, dipeptide composition, residue entropy, repeats, distribution of amino acids. In order to identify relevant features, we adopt different techniques to remove useless features or descriptors. All descriptors having low variance has been removed as they are not suitable for classification. Highly correlated or redundant has been removed to decrease the noise. Finally algorithms in Scikit-learn has been used for selecting important descriptors for developing prediction models. We employed different machine learning algorithms using python-library-scikit-learn. We implemented widely used machine learning classifiers, like KNN, DT, RF, GNB, LR, SVC and XGB [30]. In order to our models we used internal and external validations [31] [32]. The result of the generated model was analysed using various parameters called as threshold-dependent parameters and threshold-independent parameters [33] [34].We also validated the sturdiness of our model by cross checking the resistance of 22 ceftazidime resistance beta-lactamases downloaded from RGI database. Our model correctly predicted 19 ceftazidime resistance strains out of 22. We hold an opinion that this method will be very helpful in prior prediction of ceftazidime resistance/susceptibility towards any newly emerging strain of beta-lactamases. This also open vista for researchers to look for alternative therapeutic options to fight continuously emerging beta-lactamases. The method also has a major utility in doing prediction of sensitive beta-lactamase strains in metagenomics data.

## Conclusion

In conclusion, this is the first study of resistance/sensitivity prediction model development using one particular antibiotic. The study brings about in-silico model to predict resistance/susceptibility of ceftazidime antibiotic towards beta-lactamases(http://webs.iiitd.edu.in/raghava/abcrpred/). This will help in identification of ceftazidime sensitive beta-lactamases strains. Prediction can be done even when only protein sequence of any beta-lactamase is known. We believe in future, researchers will build similar model for other antibiotics. Prior prediction of sensitive antibiotics against a bacterial infection will lead to era of strain-specific antibiotics; basically, end of present hit and trial era. This will reduce time and cost of treatment as well a significant reduction in side-effects due to the treatment by inappropriate antibiotics.

## Author Contributions

LM, SSU, AD, and SP collected and processed the datasets. LM, AD, SP, SSU and GPSR implemented the algorithms. SP and AD developed the prediction models. LM, AD, SP, SSU and GPSR analysed the results. SP, NS and AD created the back-end of the web server and front-end user interface. LM, AD, SP, SSU, NS and GPSR penned the manuscript. GPSR conceived and coordinated the project and gave overall supervision to the project. All authors have read and approved the final manuscript.

## Acknowledgement

Authors are thankful to J.C. Bose National Fellowship, Department of Science and Technology (DST), Government of India, and DST-INSPIRE, NPDF-SERB, and DBT for fellowships and the financial support.

## Data Availability Statement

All the datasets generated for this study are either included in this article/Supplementary material or available at the “ABCRpred” webserver, https://webs.iiitd.edu.in/raghava/abcrpred/download.php as mentioned in the Materials and Methods section.

